# Heterologous expression of *Dictyostelium discoideum* NE81 in mouse embryo fibroblasts reveals conserved mechanoprotective roles of lamins

**DOI:** 10.1101/2023.05.31.543154

**Authors:** Jacob Odell, Ralph Gräf, Jan Lammerding

## Abstract

Lamins are nuclear intermediate filament proteins that are ubiquitously found in metazoan cells, where they contribute to nuclear morphology, stability, and gene expression. Lamin-like sequences have recently been identified in distantly related eukaryotes, but it remains unclear if these proteins share conserved functions with the lamins found in metazoans. Here, we investigate conserved features between metazoan and amoebozoan lamins using a genetic complementation system to express the *Dictyostelium discoideum* lamin-like protein NE81 in mammalian cells lacking either specific lamins or all endogenous lamins. We report that NE81 localizes to the nucleus in cells lacking Lamin A/C, and that NE81 expression improves nuclear circularity, reduces nuclear deformability, and prevents nuclear envelope rupture in these cells. However, NE81 did not completely rescue loss of Lamin A/C, and was unable to restore normal distribution of metazoan lamin interactors, such as emerin and nuclear pore complexes, which are frequently displaced in Lamin A/C deficient cells. Collectively, our results indicate that the ability of lamins to modulate the morphology and mechanical properties of nuclei may have been a feature present in the common ancestor of *Dictyostelium* and animals, whereas other, more specialized interactions may have evolved more recently in metazoan lineages.

## Introduction

Lamins are Type V intermediate filaments that form a dense protein meshwork at the inner face of the inner nuclear membrane (Aebi *et al*., 1986; Turgay *et al*., 2017). Vertebrate lamins are grouped into A-type and B-type lamins. In mammalian cells, A-type lamins include Lamins A and C (henceforth abbreviated as Lamin A/C), encoded by the *LMNA* gene and resulting from alternative splicing; B-type lamins include Lamin B1 and Lamin B2, encoded by the *LMNB1* and *LMNB2* genes, respectively. Despite the similar structures of A and B-type lamins, they have distinct functions and form separate networks in metazoan nuclei (Shimi *et al*., 2015). Lamin A/C are important modulators of nuclear shape and deformability (Sullivan *et al*., 1999; Lammerding *et al*., 2006; Swift *et al*., 2013), but also play important roles in chromatin organization, gene expression, and anchoring inner nuclear membrane proteins such as emerin to the nuclear envelope (de Leeuw *et al*., 2018; Liddane and Holaska, 2021; Kalukula *et al*., 2022). Several human diseases result from mutations in lamins or lamin-associated proteins, including various muscular dystrophies, dilated cardiomyopathies, and Hutchinson-Gilford progeria syndrome, which are collectively called laminopathies or nuclear envelopathies (Worman, 2012). These diseases are often characterized by deficiencies in mechanically active tissues such as skeletal and cardiac muscle, and cells from laminopathy disease models often have characteristic nuclear defects that include increased sensitivity to mechanical stress, disruptions in nuclear shape, and mislocalization of lamina-associated proteins (Sullivan *et al*., 1999; Swift *et al*., 2013; Cho *et al*., 2019; Earle *et al*., 2020).

Traditionally, lamins have been considered unique to metazoans, in part because of their essential functions in protecting multicellular systems from mechanical stress (Gruenbaum and Foisner, 2015). All metazoan species appear to have genes encoding for at least one lamin isoform, and in the vertebrate lineages lamins have expanded into the A- and B-types relevant for disease in humans (Peter and Stick, 2015). In the past decade, however, a number of genes that resemble the sequence of animal lamins have been found in non-metazoan lineages. For example, lamin-like sequences have been identified in choanoflagellates (*Monosiga brevicollis*) and mesomycetozoea (*Capsasproa owczaraki*), which are outgroups of metazoa and are intermediary between metazoa and fungi (Kollmar, 2015). Lamin-like sequences have also been identified in distantly related Amoebozoa (*Dictyostelium discoideum*) and Stramenopiles (*Phytophthora ramorum*), raising the possibility that lamins may have been present in the last eukaryotic common ancestor (LECA) (Gräf *et al*., 2015; Kollmar, 2015). Several of these proteins have been shown to correctly localize to the nucleus of mammalian cells and form structures that resemble lamin filaments *in vivo* (Koreny and Field, 2016). Nonetheless, functional similarities between these distantly related proteins and the animal lamins remain largely unexplored.

To investigate this question, we probed for conserved functional properties between Lamin A/C in mammals and the lamin-like protein NE81 in the amoeba *Dictyostelium discoideum*. NE81 is the most extensively characterized lamin-like protein to date, and was the first identified outside of metazoa (Krüger *et al*., 2012). NE81 has conserved sequence features with the animal lamins, including a coiled-coil rod domain, nuclear localization signal, conserved CDK1 phosphorylation site, and a C-terminal CaaX Box motif (Krüger *et al*., 2012) (Fig. 1A). These domains appear in the same pattern as in the animal lamins, and *in vitro* filament assembly experiments revealed that NE81 can form filaments of similar thickness to animal intermediate filaments (Grafe *et al*., 2019). NE81 was also found to interact with *Dictyostelium* nuclear envelope proteins, including Sun1 (a direct homolog of *SUN1* in humans) (Batsios *et al*., 2016b) and Src1 (a member of the HeH protein family that includes emerin) (Batsios *et al*., 2016a), suggesting that some of the interactions that the mammalian lamins possess may have an ancient origin. Overexpression of GFP-tagged NE81 lacking the CaaX box in *Dictyostelium* resulted in cells with increased sensitivity to shear stress, suggesting that NE81 might be involved in providing structural support to the nucleus (Krüger *et al*., 2012). However, these studies were limited by the fact that deletion of NE81 from *Dictyostelium* is lethal, preventing characterization of the mechanical properties of cells lacking NE81 at the single cell level. Furthermore, it remains unclear how NE81 functionally compares to mammalian lamins, and whether the important mechanoprotective features of vertebrate lamins were already present in a common ancestor and conserved in *Dictyostelium*, or if they are novel inventions of metazoan lineages.

**Figure 1:**
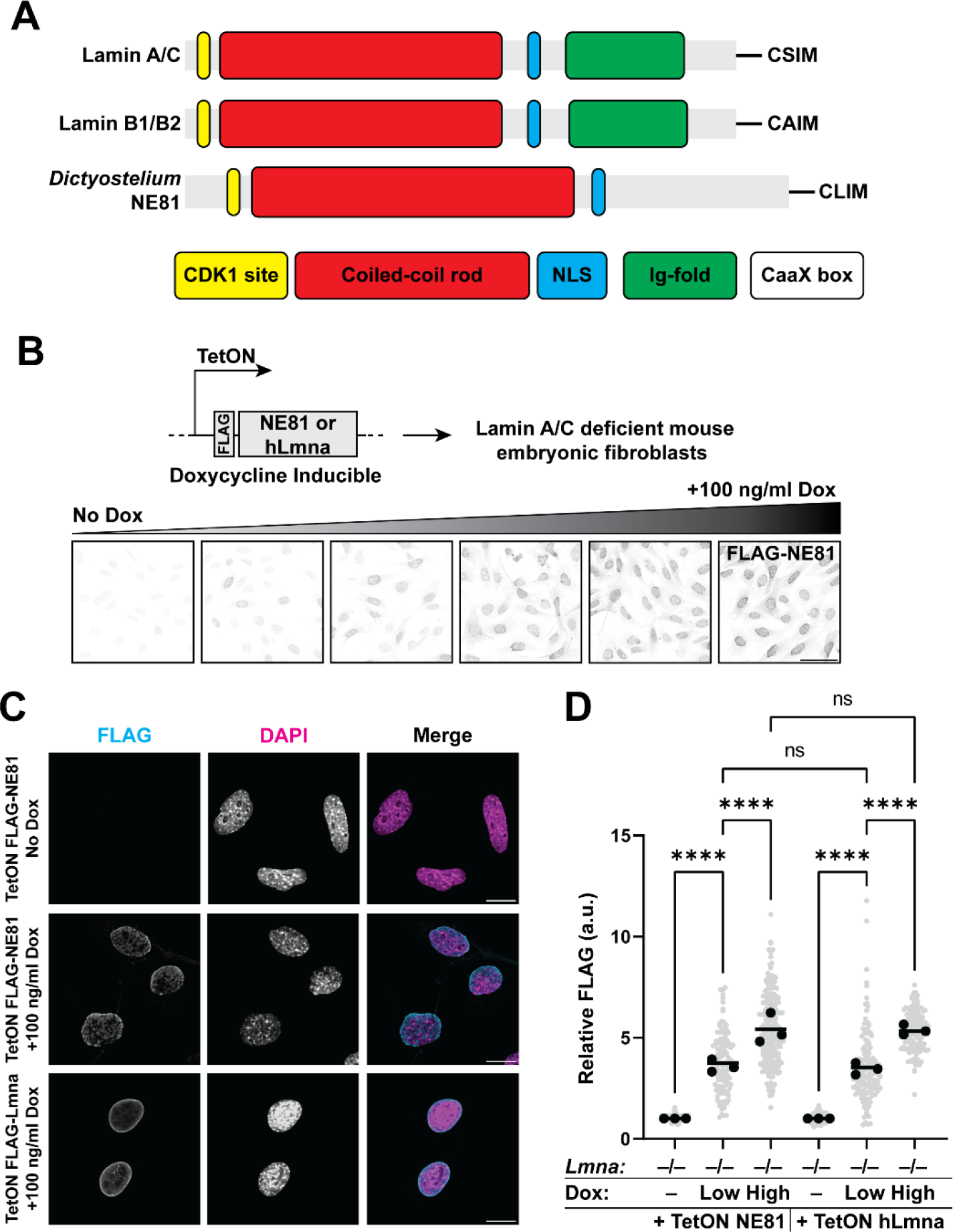
Expression of *Dictyostelium* NE81 in Mouse Fibroblasts lacking Lamin A/C. (**A**) Comparison of the major domains in human lamins versus *Dictyostelium* NE81. CDK1, cyclin-dependent kinase 1 phosphorylation site; NLS, nuclear localization signal; CaaX box is a motif recognized for post-translational modification to add a farnesyl lipid anchor, where C is a cysteine, a is any aliphatic amino acid, and X is any amino acid. (**B**) Overview and validation of the inducible constructs used in the subsequent experiments. FLAG-NE81 or FLAG-human Lamin A was cloned downstream of a doxycycline-inducible (TetON) promoter and stably inserted into *Lmna*^−/−^ MEFs to evaluate its rescue ability. Titration of doxycycline leads to concomitant increase in NE81 expression, visualized by immunofluorescence staining for the FLAG tag. Representative images shown for FLAG-NE81. Similar results were observed for FLAG-Lamin A, as quantified in panel D and Suppl. Fig. 1. (**C**) Representative confocal cross sections of the cells in (**B**) reveal that NE81 localizes to the nucleus and is enriched at the periphery, to a lesser extent than human Lamin A. Scale bar: 20 μm. (**D**) Relative expression levels of FLAG-tagged NE81 and Lamin A for each cell line and condition. Cells were immunofluorescently labeled against the FLAG tag, and relative expression levels were calculated by measuring the mean immunofluorescence intensity for each cell and normalizing values to the “no dox” condition for each replicate. “Low dox” = 12.5 ng/ml and “High dox” = 100 ng/ml. Grey dots represent measurements from individual cells, black points are means from 3 independent experiments, bar indicate overall means. ****, *p* < 0.001; ns, not significant. 150-200 cells were scored for each condition. One-way ANOVA with Tukey’s multiple comparison test.

To explore this question, we designed a genetic complementation system to ectopically express *Dictyostelium* NE81 at well-defined levels in mouse embryo fibroblasts (MEFs) lacking one or more of their endogenous lamins, and to determine to what extent NE81 could rescue loss of endogenous lamins in these cells, thereby suggesting conserved functions. The Lamin A/C deficient (*Lmna*^−/−^) and ‘triple lamin knockout’ (TKO: *Lmna*^−/−^, *Lmnb1*^−/−^, *Lmnb2*^−/−^) cell lines have been well characterized previously in terms of their mechanosensitive defects (Sullivan *et al*., 1999; Lammerding *et al*., 2004, 2006; Zwerger *et al*., 2013; Chen *et al*., 2018). Our analysis revealed that NE81 can improve nuclear circularity, provide resistance to deformation, and protect nuclear envelope integrity in *Lmna*^−/−^ and TKO MEFs. These results suggest that the ability of lamins to modulate the morphological and mechanical properties of nuclei is conserved in *Dictyostelium* and may have been one of the first functions of lamins to evolve. Importantly, however, NE81 was not able to fully compensate for the loss of mammalian lamins. In particular, NE81 was unable to rescue the localization of the nuclear envelope protein emerin or nuclear pores, suggesting that these interactions may have evolved more recently and might be restricted to metazoan lamins.

## Results

### Inducible expression of NE81 in Lamin A/C deficient mouse fibroblasts

Lamin A is a major contributor to nuclear deformability in mammalian cells, and we previously demonstrated that exogenous expression of human Lamin A can rescue impaired nuclear structure and mechanics in *Lmna*^−/−^ MEFs and myoblasts (Lammerding *et al*., 2006; Zwerger *et al*., 2013; Earle *et al*., 2020). To investigate conserved properties between mammalian lamins and NE81, we expressed either NE81 or Lamin A in *Lmna*^−/−^ MEFs and assessed to what degree NE81 can compensate for loss of endogenous lamins compared to ectopically expressed human Lamin A. An accurate interpretation of these results requires comparing cells with similar expression levels of the proteins. To enable fine control over the timing and dosage of NE81 and Lamin A levels, we cloned expression constructs for either FLAG-tagged NE81 or FLAG-human Lamin A into a doxycycline-inducible system and stably inserted the resulting expression systems into *Lmna*^−/−^ MEFs (Figure 1B-C). Titrating the doxycycline levels led to a concomitant increase in the levels of protein expression, detectable by the FLAG tag on the N-terminus of NE81 and Lamin A (Fig. 1B). Note that due to potential antibody specificity differences in recognizing the endogenous mouse versus the ectopically expressed human lamin A epitopes, we cannot precisely quantify the degree of Lamin A expression relative to wild-type cells, but estimate that both the “Low” and “High” expression results in an overexpression compared to wild-type levels (Suppl. Fig. 3B). When expressed in *Lmna*^−/−^ MEFs, NE81 localized to the nuclear periphery (Fig. 1C), as expected for a lamin protein and consistent with previous results transiently transfecting NE81 in HeLa cells (Krüger *et al*., 2012). Based on the titration experiments (Fig. 1B, D), we determined doxycycline concentrations for each construct that led to similar protein levels of NE81 and Lamin A in the cells (Fig. 1D and Suppl. Fig. S1), allowing us to perform side-by-side comparisons of the rescue ability of these proteins.

### Dosage dependent NE81 improvement of nuclear circularity in *Lmna*^*–/–*^ MEFs

After validating the expression system and identifying conditions that lead to equivalent expression levels, we compared nuclear morphology in *Lmna*^−/−^ MEFs modified to express similar levels of either NE81 or Lamin A. As expected, increasing expression of human Lamin A led to progressively more circular nuclei (Fig. 2A). In contrast, NE81 showed a biphasic effect on nuclear shape. Low levels of NE81 expression in *Lmna*^*–/–*^ MEFs improved nuclear circularity, whereas high expression levels failed to improve nuclear shape (Fig. 2A). At the high expression levels, NE81 frequently accumulated at the nuclear periphery and displaced the endogenous Lamin B1 from the nuclear envelope (Fig 2B, arrowheads). Similar displacement of Lamin B1 was observed following transient overexpression of GFP-tagged NE81 in *Lmna*^−/−^ MEFs (Fig 2C), suggesting that high levels of NE81 can displace the endogenous Lamin B1 across different expression constructs. Loss of Lamin A/C has been previously shown to partially mislocalize Lamin B1, often resulting in displacement from one pole of the nucleus (Sullivan *et al*., 1999). Quantification of Lamin B1 mislocalization confirmed that high levels of NE81 led to increased Lamin B1 mislocalization (Fig. 2D). This behavior differed from the exogenous expression of Lamin A, for which increased expression further improved Lamin B1 localization (Fig. 2D). Colocalization analysis showed that the FLAG-Lamin A signal completely overlapped with the Lamin B1 signal, whereas FLAG-NE81 had only partial colocalization with Lamin B1 (Fig. 2E). These data suggest that the biphasic, dosage-dependent effect of NE81 expression on improving nuclear circularity in *Lmna*^−/−^ MEFs is due to the propensity of NE81 to accumulate at the nuclear envelope and displace endogenous Lamin B1 when expressed at high levels, which results in aberrant nuclear morphologies at high levels of NE81 induction, offsetting the positive effect of NE81 on nuclear circularity seen at low expression levels. Because of this dosage dependent response, we used two doxycycline concentrations (“Low dox” and “High dox”) for the remainder of the experiments expressing NE81 or Lamin A in *Lmna*^−/−^ MEFs. These concentrations of doxycycline result in approximately equal levels of exogenous NE81 and human Lamin A expression at each concentration, allowing for direct comparison of the rescue potential of each construct (Figure 1D).

**Figure 2:**
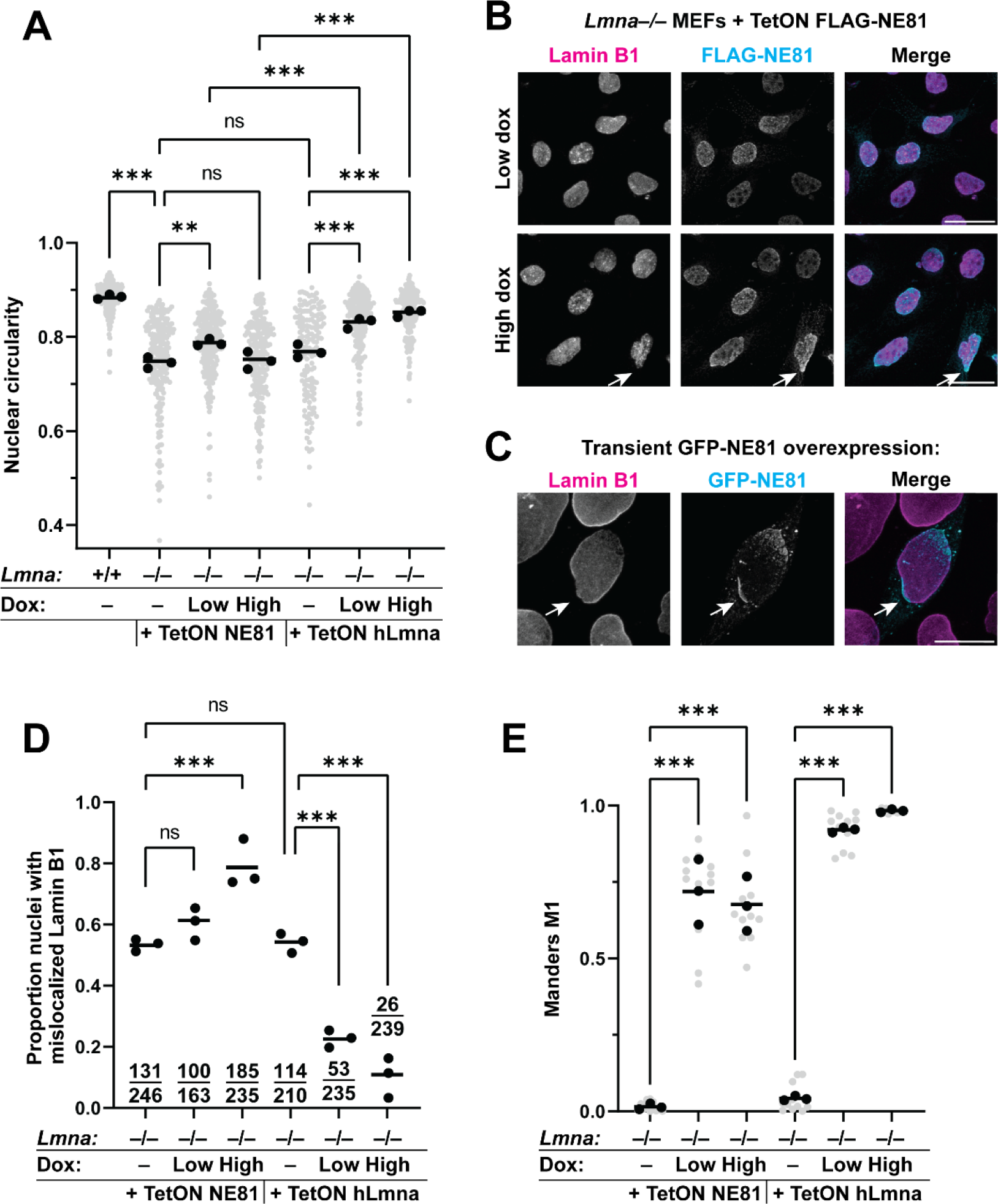
Dosage dependent NE81 rescue of nuclear circularity in *Lmna*^−/−^ MEFs. (**A**) Nuclear circularity analysis of doxycycline-inducible NE81 or human Lamin A using the doxycycline conditions identified in Figure 1. (**B**) Representative immunofluorescence images of *Lmna*^−/−^ MEFs expressing NE81 at two different expression levels reveal displacement of Lamin B1 from regions of the nuclear envelope enriched in NE81 when NE81 is expressed at high levels. Scale bar: 50 μm. (**C**) Representative images of cells with transient overexpression of GFP-NE81, showing displacement of Lamin B1 from the nuclear envelope (arrow) at regions enriched in NE81. Scale bar: 20 μm. (**D**) The proportion of nuclei with mislocalized Lamin B1 was scored by observers blinded to genotype/treatment. Points represent mean proportions from three independent experiments, bars represent the overall proportion, and counts of cells with abnormal/normal Lamin B1 are displayed before each set of points. **, *p* < 0.01; ***, *p* < 0.001; ns, not significant. Fisher’s exact test comparing proportions of abnormal/normal nuclei. (**E**) Colocalization analysis was performed using the FIJI plugin JACoP. Mander’s M1 represents the proportion of pixels in the FLAG-NE81 or FLAG-Lamin A channel that overlap with pixels in the Lamin B1 channel, with a maximum value of 1 for perfect correlation. ***, *p* < 0.001 One-way ANOVA with Tukey’s multiple comparison test. 150-200 cells were scored for each condition.

### NE81 reduces nuclear deformability in *Lmna*^*–/–*^ MEFs

Nuclei of cells with mutations in Lamin A or deletions of the *Lmna* gene are more deformable, as measured by cell stretching (Lammerding *et al*., 2006), micropipette aspiration (Pajerowski *et al*., 2007; Davidson *et al*., 2014, 2019; Earle *et al*., 2020; Bell *et al*., 2022), or atomic force microscopy (Borin *et al*., 2020). To probe differences in the mechanical properties of nuclei, we used a recently developed microfluidic micropipette aspiration assay (Davidson *et al*., 2019) outlined in Figure 3A-B. This assay enables higher throughput measurements than traditional micropipette aspiration by probing up to 18 nuclei simultaneously. In addition, micropipette aspiration induces larger nuclear deformations than cell stretching or indentation approaches, and the resistance to these large deformations has been shown previously to be governed primarily by Lamin A/C levels, whereas resistance to smaller deformations is strongly dependent on chromatin organization (Stephens *et al*., 2017). As demonstrated previously (Davidson *et al*., 2019), *Lmna*^−/−^ MEFs had substantially more deformable nuclei than wild-type MEFs (Fig. 3C-E). Exogenous expression of Lamin A and, to lesser extent, NE81, significantly reduced nuclear deformation of *Lmna*^−/−^ MEFs (Fig. 3C-E). This rescue scaled with protein expression levels; at the higher concentrations of doxycycline, nuclei were less deformable due to the higher levels of NE81 or Lamin A in the cells. However, neither NE81 nor human Lamin A was able to fully restore the wild-type level of nuclear deformability. The incomplete rescue upon expression of human Lamin A may be partly explained by the fact that these cells still lack Lamin C, and that subtle differences in the mouse versus human Lamin A sequence might hinder the rescue potential. Furthermore, both NE81 and Lamin A were tagged with an N-terminal FLAG tag, which could also affect the assembly of these proteins. As a control for contributions of the actin cytoskeleton to cell/nuclear deformability, we performed similar experiments on cells treated with Cytochalasin D, which disrupts actin filaments. Similar to what we had observed in untreated cells (Fig. 3E), NE81 significantly reduced nuclear deformability in the Cytochalasin D treated cells (Suppl. Fig. 2A-B), indicating that the observed effect is nucleus-intrinsic. To ensure that the effect on nuclear stiffness was specific to NE81 and Lamin A, and not simply because of expression of a bulk protein in the nucleus, we generated and expressed an inducible FLAG-hLmnB1 construct analogous to the constructs for NE81 and hLamin A. This construct results in overexpression of Lamin B1 relative to wild-type cells, but due to potential difference in antibody epitope recognition between the endogenous mouse Lamin B1 and exogenous human Lamin B1, we are unable to precisely quantify the degree of overexpression. Immunoblot analysis against the FLAG-tag, which is identical in all exogenous expression constructs, revealed that FLAG-hLaminB1 is expressed at levels comparable to the “High” level of Lamin A and NE81 expression described previously (Suppl. Fig. 3B). Overexpression of Lamin B1 in *Lmna*^−/−^ MEFs using this construct did not reduce nuclear deformability of *Lmna*^−/−^ MEFs (Suppl. Fig. 3), consistent with previous findings that A-type lamins are the major contributors to nuclear mechanics and that overexpression of Lamin B1 does not increase nuclear stiffness in *Lmna*^−/−^ MEFs (Lammerding *et al*., 2006). Our results demonstrate that in addition to rescuing nuclear circularity, NE81 can partially rescue nuclear deformability in *Lmna*^−/−^ MEFs when expressed at low levels, suggesting that the ability to modulate the mechanical properties of nuclei may be conserved between metazoan lamins and *Dictyostelium* NE81.

**Figure 3:**
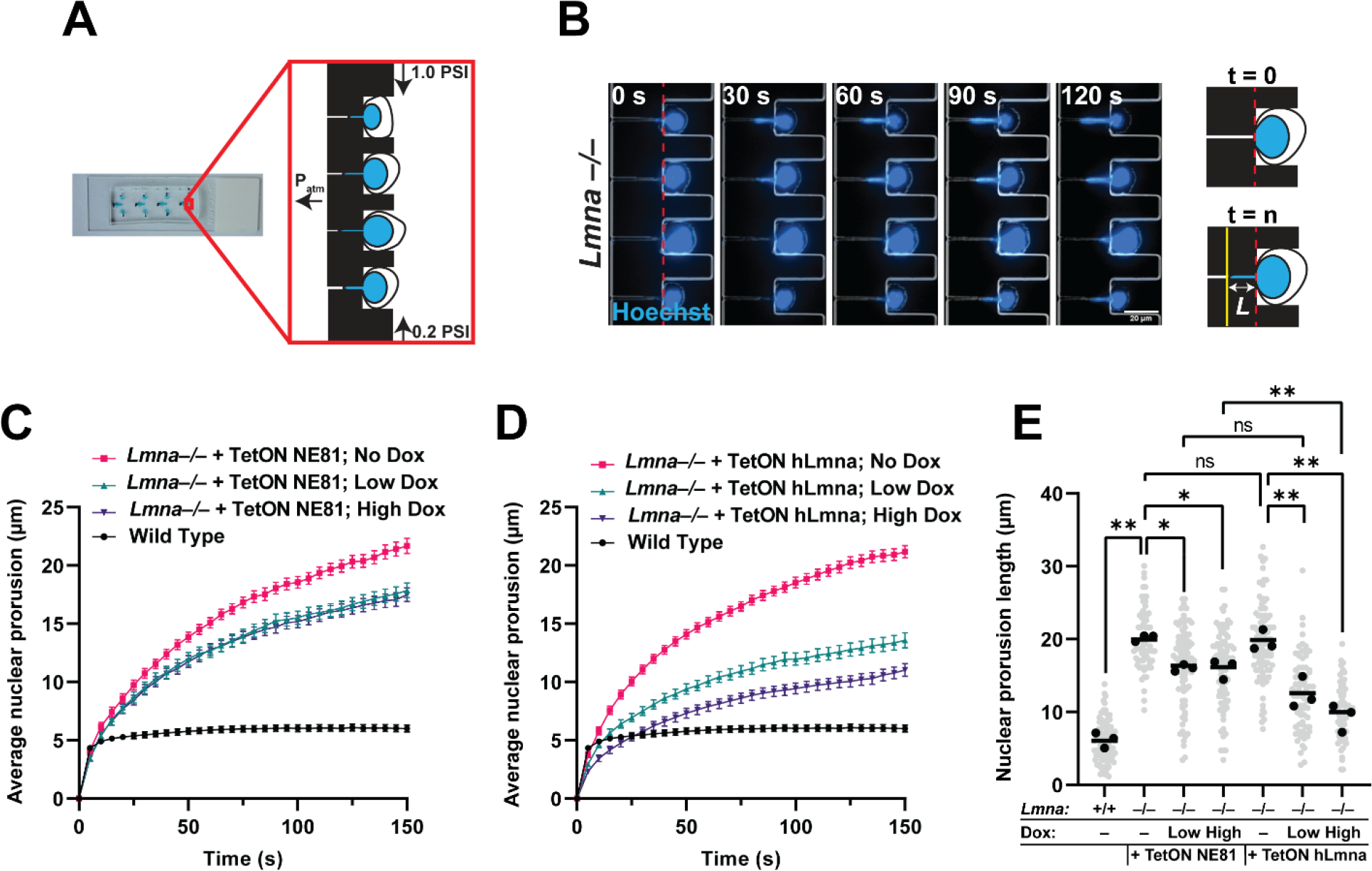
Expression of NE81 reduces nuclear deformability in *Lmna*^*−/−*^ MEFs. (**A**) Schematic of microfluidic device used for aspiration. Adapted from (Davidson *et al*., 2019). (**B**) Representative time-lapse image series of *Lmna*^*–/–*^ MEF aspiration reveals progressive increase in nuclear protrusion length (*L*) over time. (**C-D**) Quantification of average nuclear protrusion over time, with *t* = 0 being the first frame when cells entered the micropipette channel, for *Lmna*^*–/–*^ MEFs expressing inducible NE81 (**C**) or human Lamin A (**D**). Error bars represent mean ± SEM. (**E**) Quantification of nuclear protrusion lengths at *t* = 120 s reveal significant decreases in protrusion length upon dox-induced expression of NE81 or Lamin A compared to the corresponding no-dox condition. Cells were treated with “Low dox” (12.5 ng/ml), “High dox” (100 ng/ml), for 24 hours prior to aspiration. * *p*, < 0.05; ** *p*, < 0.001; ns, not significant. One-way ANOVA with Tukey’s multiple comparison test. 60-80 cells were scored for each condition/treatment.

### NE81 reduces spontaneous nuclear envelope rupture in *Lmna*^*–/–*^ MEFs

Besides increasing nuclear deformability, loss of Lamin A also increases the mechanical fragility of nuclei and increases their susceptibility to nuclear envelope rupture (Devos *et al*., 2014; Raab *et al*., 2016; Cho *et al*., 2019; Earle *et al*., 2020). To compare the effect of NE81 and human Lamin A in preventing nuclear envelope rupture in *Lmna*^−/−^ MEFs, we modified these cells with a previously established reporter for nuclear envelope rupture, cGAS-mCherry (Denais *et al*., 2016; Raab *et al*., 2016; Earle *et al*., 2020). The DNA binding protein Cyclic GMP–AMP synthase (cGAS) is normally distributed in the cytoplasm, but accumulates at sites of nuclear envelope rupture, where chromatin is exposed to the cytoplasm (Fig. 4A-B). The cGAS-mCherry reporter contains two mutations (E225A/D227A) that abolish its enzymatic activity and interferon production while still allowing binding to DNA (Denais *et al*., 2016). In cells cultured on rigid substrates such as glass, the force of apical stress fibers pushing down on the nucleus are sufficient to induce spontaneous nuclear envelope rupture, which becomes more prevalent in lamin deficient cells (Devos *et al*., 2014; Denais *et al*., 2016; Hatch and Hetzer, 2016). As expected, expression of Lamin A significantly reduced the rates of spontaneous nuclear envelope rupture in *Lmna*^−/−^ MEFs. Similarly, expression of NE81 significantly reduced the proportion of cells with cGAS-mCherry foci in *Lmna*^−/−^ MEFs, indicating reduced nuclear envelope rupture (Fig. 4C). We did not detect a significant difference between the rescue achieved with either NE81 or Lamin A when expressed at the same levels, which contrasts with the partial rescue observed in the circularity and deformability experiments. *Lmna*^−/−^ MEFs expressing either NE81 or Lamin A did not exhibit any statistically significant differences in nuclear cross-sectional area (Suppl. Fig 4), suggesting that the apical mechanical stress acting on these nuclei are similar, and differences in nuclear envelope rupture are caused by differences in nuclear mechanical stability. These results suggest that NE81, like Lamin A, may act to prevent nuclear envelope rupture by maintaining nuclear integrity, and that NE81 seems to be equally effective in preventing spontaneous nuclear envelope ruptures in *Lmna*^−/−^ MEFs.

**Figure 4:**
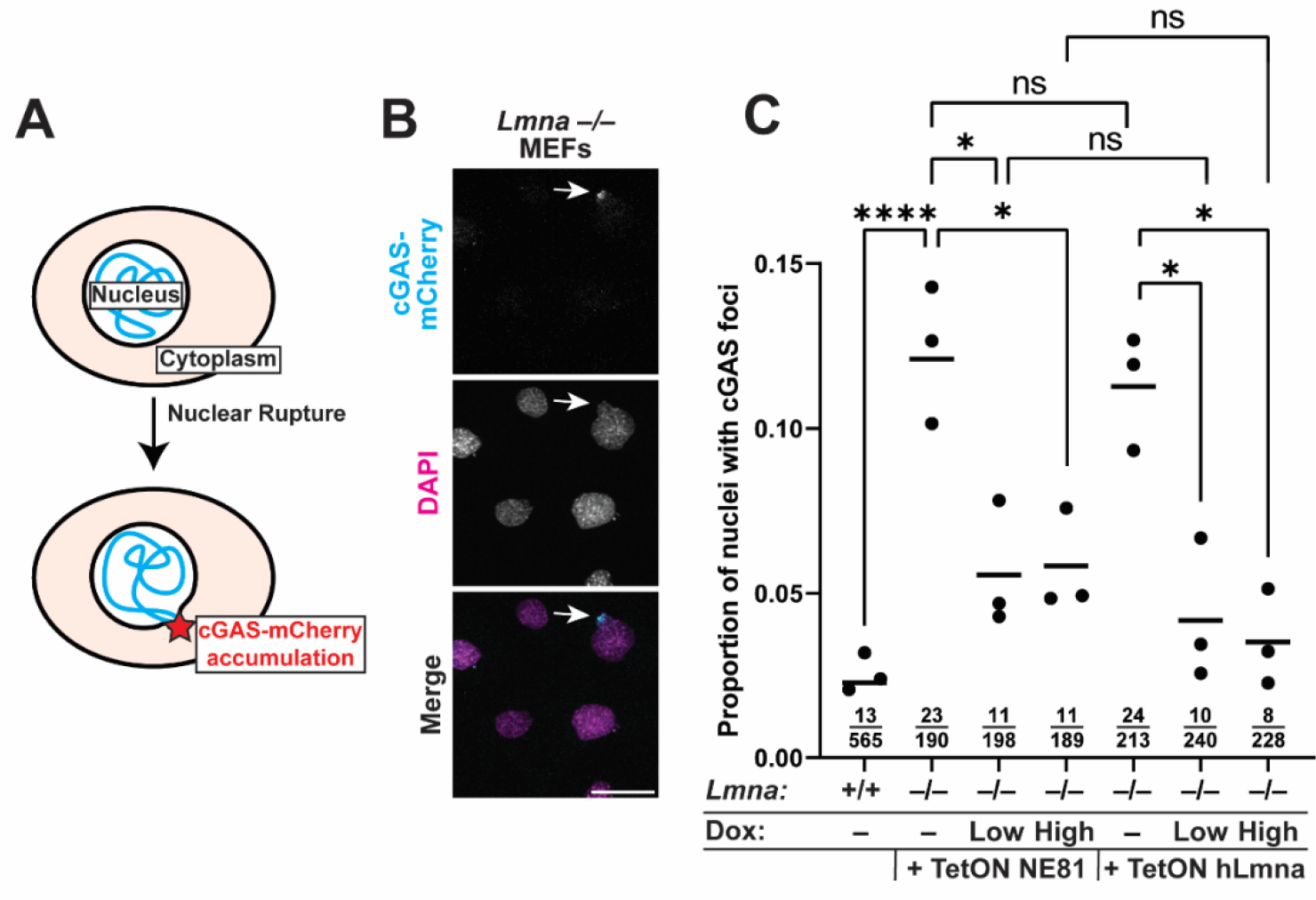
NE81 expression decreases rates of spontaneous nuclear envelope rupture. (**A**) Schematic of cGAS-mCherry reporter. The cGAS-mCherry reporter is normally located diffusely in the cytoplasm, but rapidly accumulates at sites of nuclear envelope rupture when genomic DNA is exposed to the cytoplasm. (**B**) Representative images of *Lmna*^−/−^ MEFs with cGAS-mCherry foci as puncta at the nuclear periphery, indicating nuclear envelope rupture. Scale bar: 50 μm. (**C**) Cells were scored as having or not having nuclear cGAS-mCherry puncta by observers blinded to genotype or treatment conditions. Points represent mean proportions from at least three independent experiments, bars represent total proportions of cells with cGAS foci, and fractions of “cGAS positive” (top) / total (bottom) cells are displayed below each set of points. *, *p* < 0.01; ****, *p* < 0.001; ns, not significant; Fisher’s exact test.

### NE81 rescues defects in MEFs lacking all lamins

Given the potential competition between NE81 and B-type lamins (see Fig. 2C) and to avoid any confounding effects from the presence of other endogenous lamin proteins in the *Lmna*^−/−^ MEFs, we expressed doxycycline-inducible NE81 in TKO MEFs lacking all lamins (Chen *et al*., 2018). These cells have been independently derived and have even more severe defects in nuclear morphology and deformability than the *Lmna*^−/−^ MEFs (also compare Fig. 3C-E with Suppl. Fig. 2C-D). As in *Lmna*^−/−^ MEFs, NE81 correctly localized to the nucleus in TKO cells (Suppl. Fig. S2A). Similar to the effect observed in *Lmna*^−/−^ MEFs, expression of NE81 in the TKO cells led to (partial) functional rescue of nuclear circularity (Suppl. Fig. S2B) and nuclear deformability (Suppl. Fig. S2C-D). Unlike in the *Lmna*^−/−^ MEFs, though, we did not observe any dosage-dependent effect on the nuclear circularity rescue in the TKO cells. Instead, NE81 increased nuclear circularity at all dox concentrations (Suppl. Fig. S2E). Nonetheless, across all measurements, exogenous expression of human Lamin A provided more substantial rescue in the TKO cells than expression of NE81, similar to the findings in the *Lmna*^−/−^ MEFs, suggesting that NE81 has inherent differences in assembly or interactions compared to the metazoan lamins that limit its rescue potential for nuclear shape and stiffness.

### Expression of NE81 does not restore normal nuclear localization of lamin-interacting proteins

Loss of Lamin A/C can lead to mislocalization or perturbed organization of other nuclear envelope proteins such as emerin (Sullivan *et al*., 1999), an inner nuclear membrane protein. Emerin directly binds to Lamin A and has important roles in maintaining nuclear architecture and modulating gene expression though interactions with transcriptional regulators (Lammerding *et al*., 2005; Liddane and Holaska, 2021; Fernandez *et al*., 2022). One possible explanation for the incomplete rescue observed upon NE81 expression could be that NE81 is unable to recruit emerin to the nuclear envelope. To test this hypothesis, we assessed emerin localization in *Lmna*^−/−^ MEFs expressing either NE81 or human Lamin A, and also compared these cells to *Lmna*^−/−^ MEFs and wild-type MEFs. In wild-type cells, emerin was enriched in the nucleus, but in *Lmna*^−/−^ MEFs, emerin was mislocalized to the cytoplasm and endoplasmic reticulum (Fig. 5A-B), consistent with previous findings (Sullivan *et al*., 1999). Expression of Lamin A was sufficient to restore localization of emerin to the nuclear envelope in *Lmna*^−/−^ MEFs in a dose-dependent manner (Fig. 5A-B). In contrast, NE81 was unable to rescue the nuclear localization of emerin, even at high expression levels, suggesting that NE81 cannot recruit emerin to the nuclear envelope. The lack of an apparent interaction between NE81 and emerin is likely due to the fact that emerin and other LEM-domain containing proteins appear to have evolved after the split between Amoebozoa and Opisthkonta (containing the kingdoms Fungi and Metazoa) (Brachner and Foisner, 2011). Therefore, the ability of Lamin A to interact with emerin may be feature acquired later in evolution and restricted to metazoan lineages.

**Figure 5:**
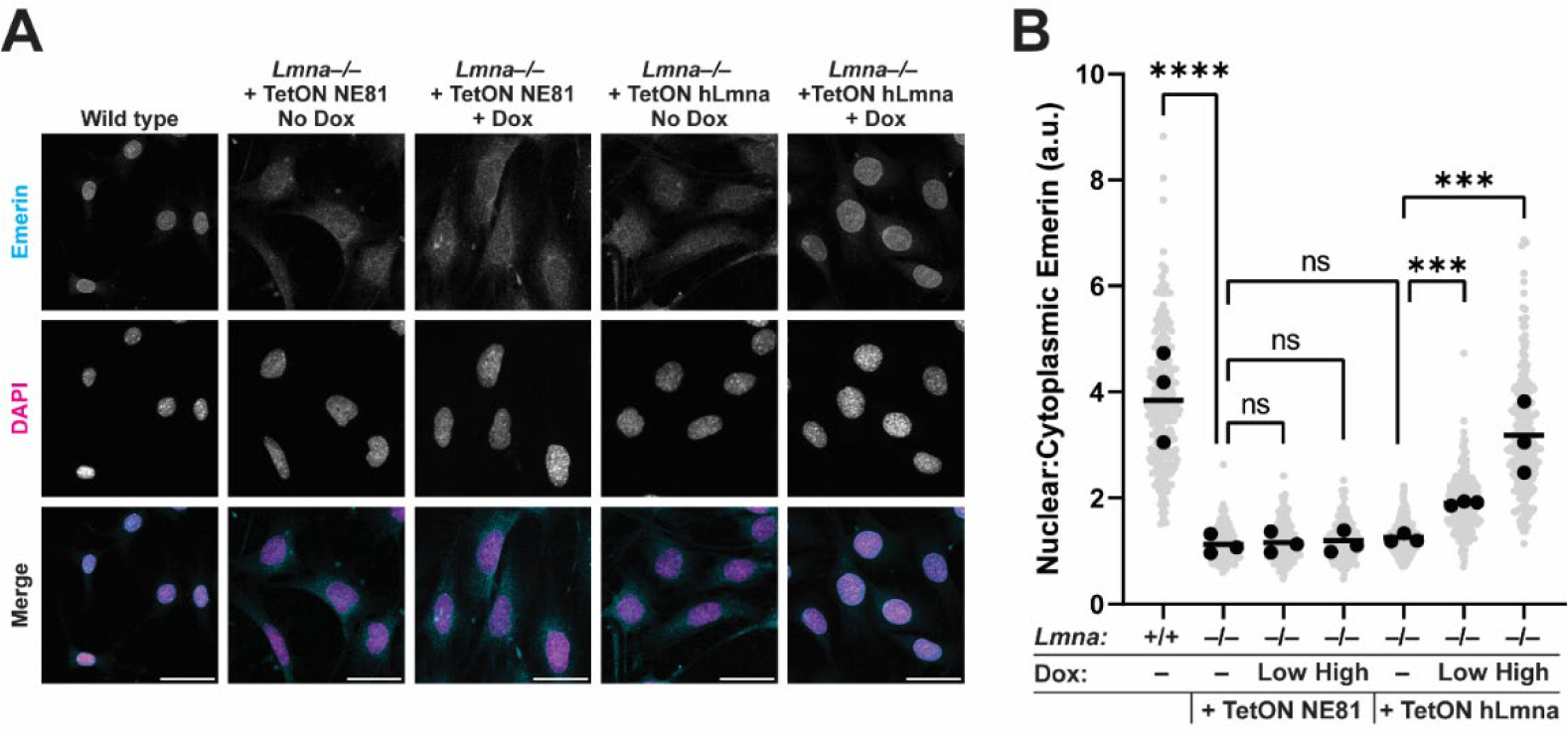
Expression of Lamin A, but not NE81, rescues emerin localization to the nuclear envelope. (**A**) Representative immunofluorescence images of cells stained for emerin reveal nuclear emerin localization in wild-type cells, and a mislocalization of emerin to the cytoplasm and ER in *Lmna*^−/−^ MEFs. Scale bar: 50 μm (**B**) Quantification of the mean nuclear/cytoplasmic ratios of emerin for individual cells. Grey points represent values calculated for individual cells, black points represent means from three independent experiments, and bars indicate the overall means. *** *p*, < 0.001; ns, not statistically significant; one-way ANOVA with Tukey’s multiple comparison test. Over 200 cells were scored per condition.

In addition to retaining inner nuclear membrane proteins at the nuclear envelope, lamins play a crucial role in positioning nuclear pore complexes (NPCs) along the nuclear envelope (Guo and Zheng, 2015; Chen *et al*., 2018). Accordingly, TKO MEFs exhibit a pronounced mislocalization of NPCs, with a clustering of NPCs to one pole or side of the nucleus (Fig. 6A), matching previous findings (Chen *et al*., 2018). To probe possible conserved interactions between NE81 and NPCs, we immunofluorescently labeled TKO cells expressing either NE81 or Lamin A with a pan-NPC antibody. As expected, expression of Lamin A restored the uniform distribution of NPCs along the nuclear periphery; in contrast, expression of NE81 did not improve the abnormal NPC distribution in TKO cells (Fig. 6A-B). These results suggest that NE81 is unable to reposition NPCs. It is possible that NE81 interacts with NPCs in TKO MEFs but cannot form sufficiently stable filaments or networks to influence their spacing. Alternatively, mammalian/*Dictyostelium* NPCs may have diverged from one another, resulting in an incompatibility of lamin binding sites between NE81 and mammalian NPCs.

**Figure 6:**
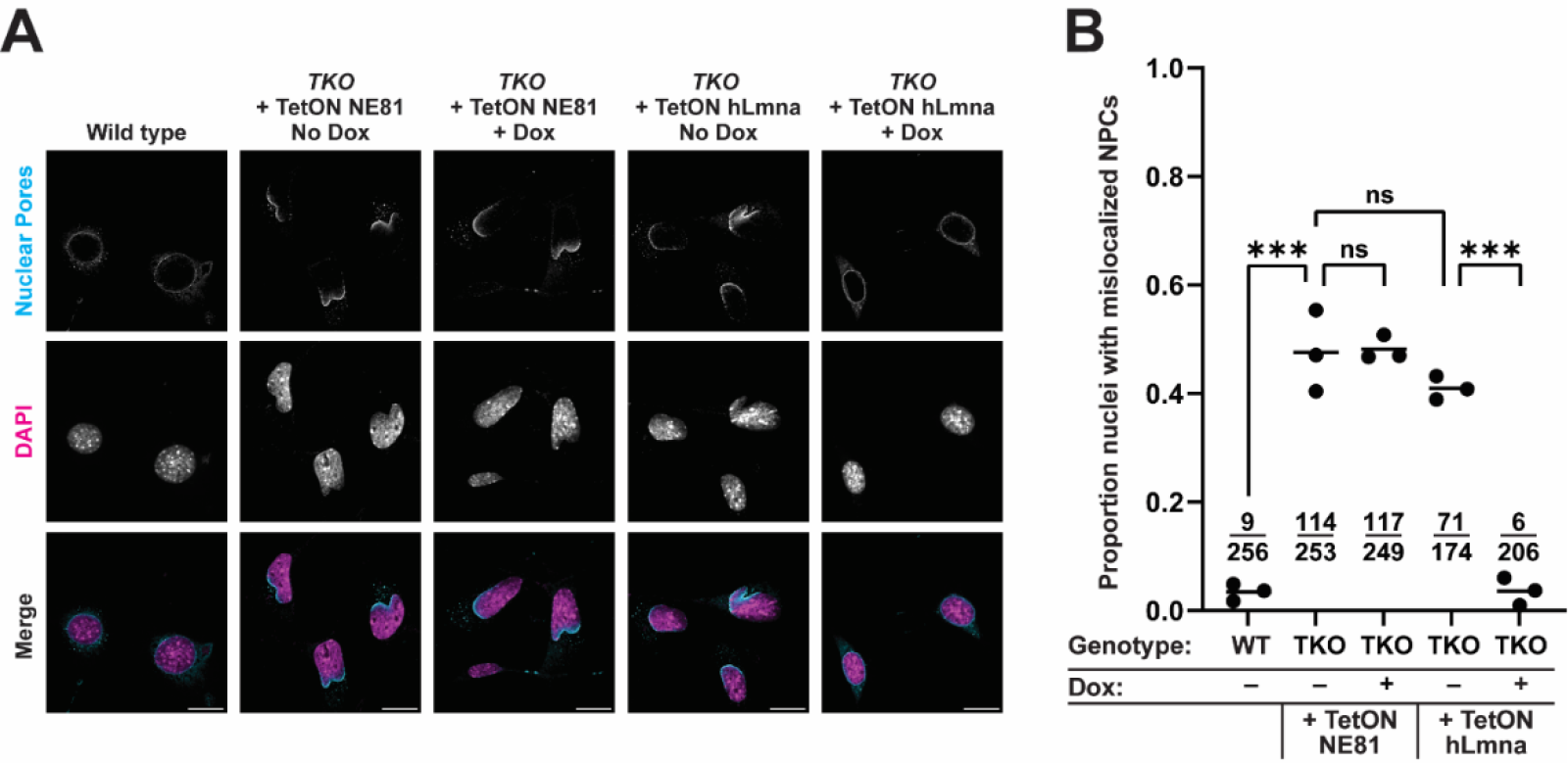
Expression of Lamin A, but not NE81, rescues nuclear pore complex (NPC) distribution. (**A**) Representative immunofluorescence images of cells stained for a pan-NPC marker (Mab414) reveals an even distribution of NPCs at the nuclear periphery in wild-type MEFs, and mislocalization and clustering of NPCs to one pole of the nucleus in cells lacking all lamins. Scale bar: 50 μm. (**B**) Quantification of the proportion of nuclei with mislocalized nuclear pores. Nuclei were scored as having “normal” or “abnormal” nuclear pore distribution at the nuclear periphery by observers blinded to genotype or treatment. Counts of “abnormal” / total nuclei are displayed below each set of points. ***, *p* < 0.001; ns, not statistically significant; Fisher’s exact test.

## Discussion

Using a genetic complementation approach with robust control over protein expression levels, we showed that the *Dictyostelium* lamin-like protein NE81 can rescue morphological and mechanical defects that arise in *Lmna*^−/−^ and TKO MEFs. Exogenous expression of human Lamin A or NE81 in cells lacking one or more endogenous lamins improved nuclear circularity, reduced nuclear deformability, and reduced rates of nuclear envelope rupture when expressed, suggesting that the lamin-like protein found in the common ancestor of *Dictyostelium* and metazoans may have already been able to provide structural support to the nucleus. However, NE81 was unable to fully compensate for the loss of Lamin A/C, evidenced by the partial rescue in the nuclear shape and deformability experiments. Whereas NE81 shares some structural functions with metazoan lamins, it lacks other functions such as emerin recruitment to the inner nuclear membrane or positioning of metazoan nuclear pores. These results suggest that in metazoans, lamins have evolved to fulfil novel, diverse roles.

The diversity of lamins in vertebrates is attributed to two rounds of whole genome duplication early in vertebrate evolution, which gave rise to the A-type lamin genes (*LMNA* and *LIII*, which is present in amphibians but lost in mammals), and the B-type lamins (*LMNB1* and *LMNB2*) (Kollmar, 2015; Peter and Stick, 2015). Most invertebrates, including *C. elegans*, have only a single lamin isoform that more closely resembles the B-type lamins in terms of structure and mobility in the nuclear envelope, but retains functions of both the A-types and B-types (Bank and Gruenbaum, 2011). Extensive characterization of the mechanical properties of the *C. elegans* lamin has improved our understanding of conserved lamin functions in the common ancestor of vertebrates/invertebrates. Both Ce-lamin and mammalian lamins control nuclear shape, stiffness, and deformability, recruit emerin to the nuclear envelope, and position nuclear pores (Liu *et al*., 2000; Bank and Gruenbaum, 2011). Like Ce-lamin, NE81 is considered more similar to B-type lamins (Gräf *et al*., 2015), due to its sequence, farnesylation, and mobility at the nuclear envelope. Removal of Lamin B1 from MEFs results in little or no change to the mechanical properties of nuclei (Lammerding *et al*., 2006), while increasing the rate of nuclear envelope ruptures (Denais *et al*., 2016; Hatch and Hetzer, 2016), suggesting distinct roles for the A- and B-type lamin networks. Our data are consistent with the classification of NE81 as a B-type lamin, because NE81 expression could substantially reduce nuclear envelope rupture events while having only a partial effect on nuclear morphology and deformability. It is worth noting that whereas expression of NE81 and Lamin A reduced nuclear deformability of *Lmna*^−/−^ MEFs, expression of Lamin B1 did not. NE81 is evolutionarily more closely related to the lamins found in basal metazoans, like Ce-lamin, which also possess features attributable to the vertebrate A- and B-type lamins. Our data suggest that the lamin-like protein found in a common ancestor of *Dictyostelium* and metazoa may have already possessed functions attributed to metazoan lamins, including supporting nuclear shape and stability, albeit to a limited extent. Further experiments will need to be performed to examine the functional characteristics of other, diverse lamin-like proteins to rule out the possibility that the properties of NE81 observed in this study simply arose by convergent evolution, instead of a shared common ancestor. A greater understanding of the lamin-like proteins found in distantly related eukaryotes will shed light on the underlying diversity present in the primordial lamin, upon which natural selection may have acted in early animals and vertebrates to result in the expansion and specialization of the A-type and B-type lamins.

Given the similarities between NE81 and the B-type lamins, as well as the vast, independent evolution of the amoebozoan and metazoan lineages, it is perhaps not surprising that expression of NE81 did not fully rescue loss of Lamin A/C. In the nuclear circularity and deformability experiments, NE81 significantly reduced the defects that occurred following the loss of endogenous Lamin A/C but, compared to the rescue achieved with exogenous Lamin A, the rescue with NE81 was incomplete. This partial rescue was not due to differences in expression, as we compared cell lines with similar protein levels of NE81 and Lamin A. The partial rescue with NE81 may be due in part to an incompatibility between NE81 and the B-type lamins, as both proteins retain farnesylation at their C-terminus (Krüger *et al*., 2012), and high levels of NE81 displaced the endogenous Lamin B1 from the nuclear envelope, possibly by competing for attachment sites at the inner nuclear membrane. We hypothesize that *Lmna*^−/−^ MEFs have a baseline level of Lamin B1 mislocalization (Sullivan *et al*., 1999), making them more prone to nuclear envelope rupture, and that NE81 can anchor the nuclear membrane to the nuclear lamina in these regions, thereby reducing the rate of spontaneous nuclear envelope rupture. However, even in the TKO MEFs, we observed a difference in rescue ability between NE81 and Lamin A, despite the absence of other B-type lamins in these cells, suggesting that the rescue potential of NE81 is also limited by other differences between NE81 and lamins. Despite resembling the overall conserved lamin architecture, NE81 shares very little sequence identity with human Lamin A, and these large sequence differences may lead to deficiencies in filament formation that prevent the normal assembly of NE81 in mammalian nuclei and limit its ability to rescue mechanical or morphological defects. We also observed that exogenous expression of human Lamin A could not fully restore wild-type levels of circularity, deformability, or rupture. This may be due to small sequence differences between the human and mouse proteins, the absence of Lamin C in these cells, or the presence of the N-terminal FLAG tag, which could impair the normal assembly and filament formation of the lamina. We intentionally chose a small tag for these experiments, because previous work on NE81 has suggested that a bulky GFP tag inhibits normal protein assembly and function (Krüger *et al*., 2012; Grafe *et al*., 2019), and the FLAG tag enables us to directly compare protein levels between ectopically expressed NE81 and Lamin A. Expression of NE81 in *Lmna*^−/−^ MEFs was unable to rescue the normal distribution of emerin to the nuclear interior. Emerin is part of a family of proteins containing the LEM (LAP2 (lamina-associated polypeptide 2)/emerin/MAN1) domain, a bihelical motif that mediates binding to the chromatin binding protein BAF (barrier to autointegration factor) (Brachner and Foisner, 2011). The LEM domain is thought to have evolved from the HeH (helix-extension-helix) family, which is capable of interacting directly with DNA similar to the LEM domain (Brachner and Foisner, 2011). *Dictyostelium* has a single HeH family member, Src1, which has been indicated to interact with NE81 *in vivo* based on proximity-dependent biotinylation assays (Batsios *et al*., 2016b, 2016a). However, emerin and Src1 share low consensus, and *Dictyostelium* does not have a homolog of BAF, which appears to be restricted to metazoan lineages. It is perhaps not surprising then that expression of NE81 cannot rescue emerin localization, since *Dictyostelium* does not have an equivalent of emerin, and amoebae do not share the co-evolution that occurred between lamins, LEM domain proteins, and BAF in metazoa. Since emerin contributes to nuclear stability and morphology (Lammerding *et al*., 2005; Rowat *et al*., 2006; Guilluy *et al*., 2014), the inability of NE81 to recruit emerin to the nuclear envelope may also contribute to the incomplete rescue observed upon NE81 expression in *Lmna*^−/−^ MEFs. In *Dictyostelium*, cells lacking NE81 display abnormal chromatin organization (Krüger *et al*., 2012), suggesting a possible link between NE81 and chromatin, but further studies will be needed to determine the significance of NE81-interacting proteins such as Src1 in this interaction to investigate whether NE81 can modulate chromatin in the absence of BAF.

Expression of NE81 was also unable to rescue the distribution of nuclear pores in cells lacking all lamins. Interactions between lamins and nuclear pores are highly conserved in animals, and are present in basal metazoans including *C. elegans* (Liu *et al*., 2000). As mentioned previously, NE81 has low amino acid identity to the mammalian lamins, and it is possible that it has evolved a unique binding motif that can position *Dictyostelium* NPCs, but that is incompatible with binding metazoan NPCs. Furthermore, although NPCs are structurally well conserved between *Dictyostelium* and mammalian cells, sequence identities of the individual nuclear pore components are relatively low (Beck *et al*., 2007), which may prevent interaction between NE81 and mammalian NPCs. It is also possible that the filaments formed by NE81 are not stable enough in mammalian cells to influence the spacing of nuclear pores, or that NE81 does not assemble into sufficiently large filaments *in vivo*, i.e., inside the cell. Further experiments will be needed to determine the structure of NE81 assemblies in the cell nucleus, and if NE81 interacts with nuclear pores in *Dictyostelium*, which would suggest that the ability of lamins to interact with nuclear pores is conserved in lamin-like proteins outside of metazoa.

Altogether, our results expand the body of knowledge on lamin evolution through heterologous expression of an amoebozoan lamin-like protein in mammalian cells. By directly comparing the rescue potential of NE81 and human Lamin A, we reveal that this lamin-like protein can partially rescue nuclear circularity, reduce nuclear deformability, and prevent nuclear envelope ruptures, suggesting that these conserved features were present in a lamin-like protein found in the common ancestor of *Dictyostelium* and animals. However, NE81 cannot fully compensate for the loss of Lamin A/C and is unable to redistribute emerin or nuclear pores. These results suggest that in animals, lamins have further adapted to fulfil novel, diverse roles. This heterologous expression approach will be helpful to elucidate the specific functions of other lamin-like proteins, which could identify specific domains and features responsible for these functions and lead to a more complete understanding of lamins.

## Methods

### Cell culture and genetic modification

Wild-type and Lamin A/C deficient MEFs were a generous gift from Colin Stewart and have been extensively characterized previously (Sullivan *et al*., 1999; Lammerding *et al*., 2004, 2006). MEFs lacking Lamin A/C, B1, and B2 were provided by Stephen Young (Chen *et al*., 2018). Cells were maintained in DMEM supplemented with 10% FBS and 1% Pen strep. Transient transfection was carried out using Lipofectamine 3000 according to manufacturer’s instructions. Stable genetic manipulations were achieved using pseudovirus particles or with the Piggybac transposase system as described previously (Denais *et al*., 2016; Earle *et al*., 2020). Antibiotic selection was performed using Puromycin at 3 μg/ml and Blasticidin at 4.5 μg/ml for at least one week, until all cells in a ‘non-transformed’ control well had died. Clonal isolation was performed by serial dilution in a 96-well plate. Putative clones were expanded and screened by immunofluorescence staining for homogenous levels of protein expression. Dox titrations were performed in 24 well plates using 1:2 serial dilutions to yield progressively more dilute doxycycline conditions.

### Genetic construct information

Codon-optimized NE81, as described previously (Krüger *et al*., 2012) was kindly provided by Ralph Gräf. FLAG-NE81 was cloned using Gibson assembly into the pCDH-CMV-hLamin_A-IRES-copGFP-EF1-puro backbone (Earle *et al*., 2020) following restriction digest with EcoR1 and Not1. Codon optimized NE81 sequence was amplified from the plasmid pIS538 (Krüger *et al*., 2012) using the PCR primers 5’-AGAAGATTCTAGAGCTAGCGAATTCCACCATGGACTACAAAGACGATGACGACAAGATGGACATGAGCAAGAAAAAG-3’, which adds a FLAG tag (DYKDDDDK), and 5’-GGGGGAGGGAGAGGGGCAGCGGCCGCTTACATGATCAGACAATTTGATTTC-3’. Doxycycline-inducible NE81 was cloned using Gibson assembly into pPB-rtTA-hCas9-puro-PB (Wang *et al*., 2017) cut with Nhe1 and Age1. FLAG-NE81 was amplified from the previous plasmid using the primers 5’-ACCCTCGTAAAGGTCTAGAGACCATGGACTACAAAGAC-3’ and 5’-CCGTTTAAACTCATTACTAATTACATGATCAGACAATTTGATTTC-3’. As controls, FLAG-human Lamin A was cloned into the same backbones. For constitutive expression, FLAG-hLmna was amplified from the pCDH-CMV-hLamin_A-IRES-copGFP-EF1-puro backbone (Earle *et al*., 2020) using the PCR primers 5’-AGAAGATTCTAGAGCTAGCGCACCATGGACTACAAAGACGATGACGACAAGATGGAGACCCCGTCCCAG-3’ and 5’-GGGGGAGGGAGAGGGGCAGCTTACATGATGCTGCAGTTCTGG-3’. Doxycycline-inducible FLAG-hLmna was cloned into the pPB-rtTA-hCas9-puro-PB backbone (Wang *et al*., 2017) using Gibson assembly following amplification of FLAG-hLmna from the previous plasmid using the PCR primers 5’-ACCCTCGTAAAGGTCTAGAGACCATGGACTACAAAGAC-3’ and 5’-CCGTTTAAACTCATTACTAATTACATGATGCTGCAGTTC-3’. The FLAG-hLmnB1 construct was generated by amplifying the human Lamin B1 sequence using the PCR primers 5’– ACCCTCGTAAAGGTCTAGAGACCATGGACTACAAAGATGACGACAAGATGGCGACTGCGACCCCC-3’ and 5’-CCGTTTAAACTATTACTAACTACATAATTGCACAGCTTCTATTGGATGCTCTTG-3’ and cloning the resulting product into the pPB-rtTA-hCas9-puro backbone described above. Following cloning, all plasmid sequences were verified by Sanger sequencing of the inserts. For Piggybac transposition, plasmids containing the insert were co-transfected with a plasmid containing a hyperactive transposase (2:1 vector plasmid: hyperactive transposase plasmid) using the Purefection system according to manufacturer’s instructions.

### Immunofluorescence

Cells were seeded on Fibronectin-coated glass coverslips for at least 24 hours before fixation and staining. When cells were to be incubated with doxycycline, cells were seeded at least 12 hours in advance before the media was replaced with doxycycline-containing media. Fixation was performed with 4 % paraformaldehyde in PBS for 15 minutes, followed by permeabilization in Tween and Triton-20 three 3-minute washes. Cells were blocked in 3 % BSA for 1 hour, then primary antibodies were added for 1 hour at room temperature or overnight at 4°C. Coverslips were washed 3-times, then secondary antibodies added at 1:250 dilution using Alexa Fluor-conjugated secondary antibodies. To stain DNA and for nuclear thresholding, DAPI was added 1:1000 and incubated for 15 minutes, followed by three 5-minute washes. Coverslips were mounted on glass slides with mowiol and left to harden overnight. Imaging was performed on an inverted Zeiss Observer Z1 microscope and CCD camera (Photometrics CoolSNAP KINO) using 20× air (NA = 0.8) and 63× oil (NA = 1.4) immersion objectives. Airy units for all images were set between 1.5 and 2.5. The image acquisition for micropipette aspiration experiments was automated through ZEN (Zeiss) software.

### Image analysis. Nuclear circularity

Measurements of nuclear circularity and intensity of different fluorescently conjugated antibodies was determined using a FIJI macro. Briefly, this macro performs a background subtraction and thresholds the image based on the DAPI channel to identify nuclei, and then measures the circularity of each nucleus and mean intensity in each channel using the Analyze Particles function. This macro was also used to determine the nuclear areas presented in Suppl. Fig. 4. **Qualitative counting of cell phenotypes:** For counts of nuclei with Lamin B1 mislocalization, images were blinded and nuclei were scored as having or not having mislocalized Lamin B1. “Mislocalized lamin B1” was defined as when there was a clear absence of Lamin B1 signal at the nuclear periphery or at one or more poles of the nucleus. For counts of cGAS-mCherry positive nuclei, images were blinded and nuclei were scored as “cGAS positive” or “cGAS negative” based on the presence or absence of mCherry puncta adjacent to each nucleus. **Nuclear/cytoplasmic emerin**: For measurements of nuclear/cytoplasmic emerin, nuclear emerin was calculated as above, and for each cell, cytoplasmic emerin was obtained by manually drawing a ROI adjacent to the corresponding nucleus for each cell. Then, nuclear emerin intensity was divided by cytoplasmic emerin intensity for each cell. **Colocalization analysis**: Performed in ImageJ using the JACoP (Just Another Colocalization Plugin) plugin (Bolte and Cordelières, 2006), following manual thresholding to identify nuclear regions. In all image analysis, cells on the edges of the images, mitotic cells, apoptotic cells, or dead cells were excluded manually from the dataset. Additionally, efforts were made to only include cells seeded at approximately equal densities, and replicates with abnormal seeding densities were excluded from the analysis.

### Immunoblotting

Cells were seeded in 6-well plates and lysed using a high salt RIPA buffer. To extract lamins/NE81, lysates were vortexed for 5 minutes, sonicated 30 seconds at 36% amplitude, boiled for 2 minutes, centrifuged at 4°C for 10 minutes and stored at –80°C. Protein concentration was determined using Bradford assay. Equal amounts of protein lysates were denatured in 5× Laemmli buffer by boiling for 3 minutes, loaded onto 4-12% Bis-Tris gels, run for 1.5 hours at 100 V, then transferred for 1 hour at 16 V onto membrane. Membranes were probed with primary antibodies at room temperature for one hour or overnight at 4°C. Secondary antibodies were added for one hour, and membranes were imaged using Odyssey Licor scanner, then cropped and brightness/contrast was adjusted using Image Studio software. Immunoblot band intensities were quantified using Image Studio Lite (Ver 5.2) using the automatic band detection function.”

### Micropipette aspiration assay

Micropipette aspiration was performed according to a previously established protocol (Davidson *et al*., 2019). In brief, cells were suspended in a 2% BSA solution supplemented with 0.2% FBS and 10 mM EDTA to prevent clumping or cell adherence. Hoechst was added 1:1000 immediately before the cell suspension was transferred to devices. For the experiments described in Suppl. Fig. 2, the actin cytoskeleton was disrupted in a subset of cells by treating cells with 4 μM Cytochalasin D for 20 minutes prior to adding the cells to the micropipette devices. Cells were perfused into the devices at 1.0 psi, and counter pressure was applied to the bottom channel at 0.2 psi. Images were acquired every 5 seconds for 40 frames, and nuclear protrusion length was determined for each micropipette pocket at each frame using a custom-written MatLab script (available at: https://github.com/Lammerding/MATLAB-micropipette_analysis). Nuclear stiffness was inferred from the protrusion lengths over time.

### Statistical analysis and figure generation

All analyses were performed using GraphPad Prism. Information on statistical tests used, cell counts, and significance values are present in each figure caption. Experiments were performed a minimum of 3 independent times, and for qualitative image analysis, observers were blinded to genotype or treatment conditions when scoring phenotypes. Our statistical analysis was developed in close consultation with the Cornell Statistical Consulting Unit. Figures were assembled using Adobe Illustrator.

## Acknowledgements

We are grateful to Stephen Young and Loren Fong for providing TKO MEFs lacking Lamin A/C, B1, and B2. We thank the Biotechnology Resource Center (BRC) Flow Cytometry Facility (RRID:SCR_021740) and sequencing facility (RRID:SCR_021727) at the Cornell Institute of Biotechnology for their resources and technical assistance. This work was performed in part at the Cornell NanoScale Science & Technology Facility, a member of the National Nanotechnology Coordinated Infrastructure, which is supported by the National Science Foundation (award NNCI-2025233). This work was supported by awards from the Volkswagen Foundation (A130142 to J.L.), the National Institutes of Health (R01 HL082792, R01 GM137605 to J.L.), and the National Science Foundation (URoL-2022048 to J.L.). The content of this manuscript is solely the responsibility of the authors and does not necessarily represent the official views of the National Institutes of Health.

