## Supplemental Materials for "Heterologous expression of *Dictyostelium discoideum* NE81 in mouse embryo fibroblasts reveals conserved mechanoprotective roles of lamins"

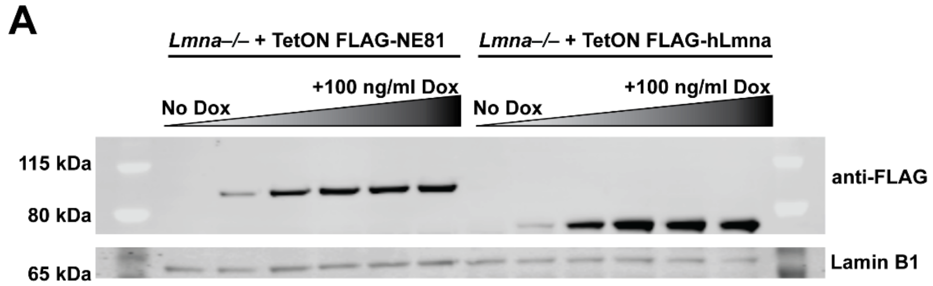


**B**

| Genotype: | Dox: | FLAG Intensity: | Lamin B1 Intensity: | FLAG/ Lamin B1: |
|---|---|---|---|---|
| *Lmna–/–* + FLAG-NE81 | – | 504 | 2850 | 0.18 |
| | Low | 25100 | 2550 | 98.43 |
| | High | 43600 | 3840 | 113.54 |
| *Lmna–/–* + FLAG-hLmnA | – | 1930 | 2470 | 0.78 |
| | Low | 29600 | 2470 | 119.84 |
| | High | 45800 | 2050 | 223.41 |

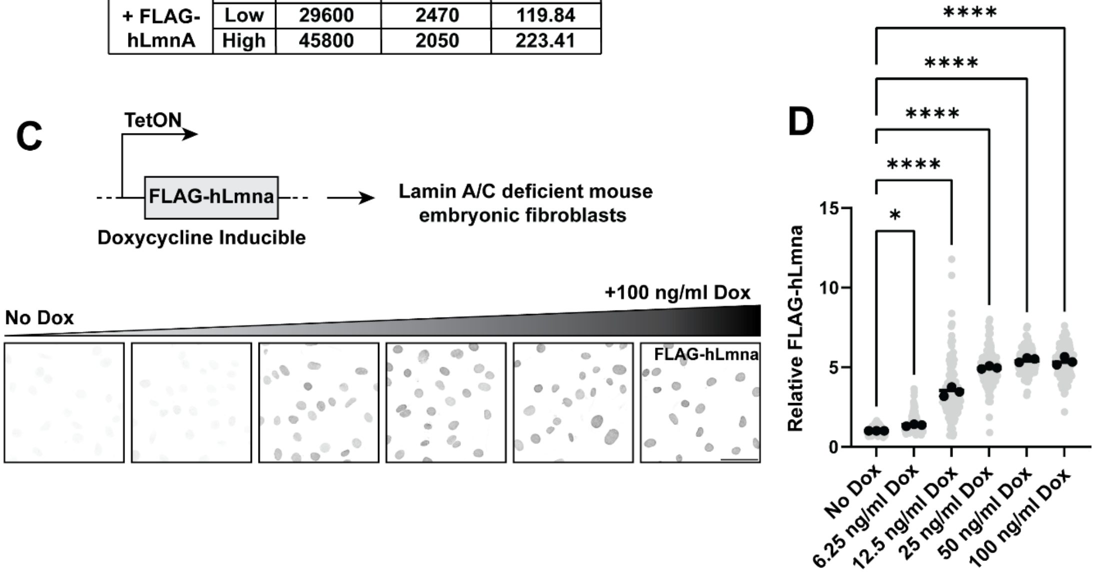


**Supplemental Figure 1: Inducible expression of NE81 or human Lamin A in Lamin A/C deficient MEFs**. (**A**) Western blot analysis on lysates collected from cells treated with increasing concentrations of doxycycline for 24 hours. Membrane was probed with anti-FLAG and anti-Lamin B1 antibodies. (**B**) Quantification of FLAG-NE81 or FLAG-hLmna band intensities from the blot depicted in (A). (**C**) Representative immunofluorescence images of cells expressing Doxycycline-inducible hLmna treated with the same dox concentrations as in (A). Scale bar: 50 µm. (**D**) Quantification of the FLAG intensity relative to the "no dox" condition for each dox concentration reveals a progressive increase in FLAG signal as the doxycycline levels increase. * $p < 0.05$; **** $p < 0.001$. One-way ANOVA with Tukey's multiple comparison test.

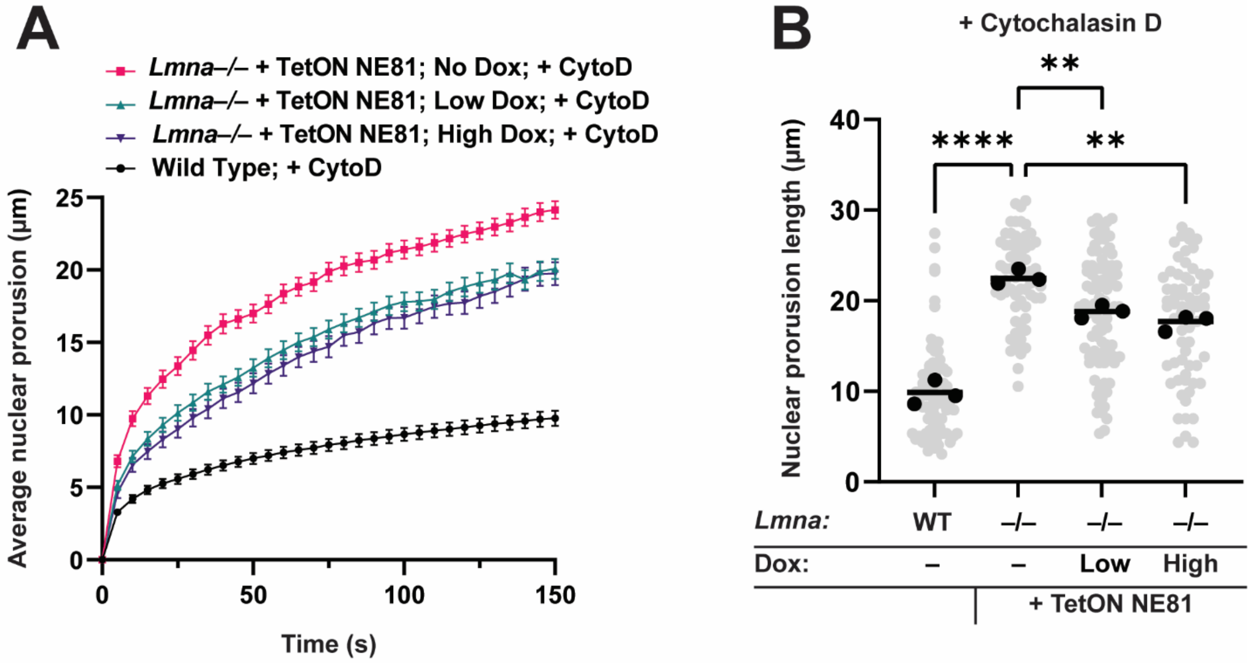


**Supplemental Figure 2: Disruption of actin cytoskeleton with Cytochalasin D.** (**A**) Quantification of average nuclear protrusion over time, as in Figure 3. All cells were treated with 4 µM CytoD for 20 minutes prior to the start of aspiration. (**B**) Quantification of nuclear protrusion lengths 120 s after cells entered micropipette pocket, as in Figure 3. ** p, < 0.01; **** p, < 0.001; One-way ANOVA with Tukey's multiple comparison test. 65-90 cells were scored for each condition/treatment.

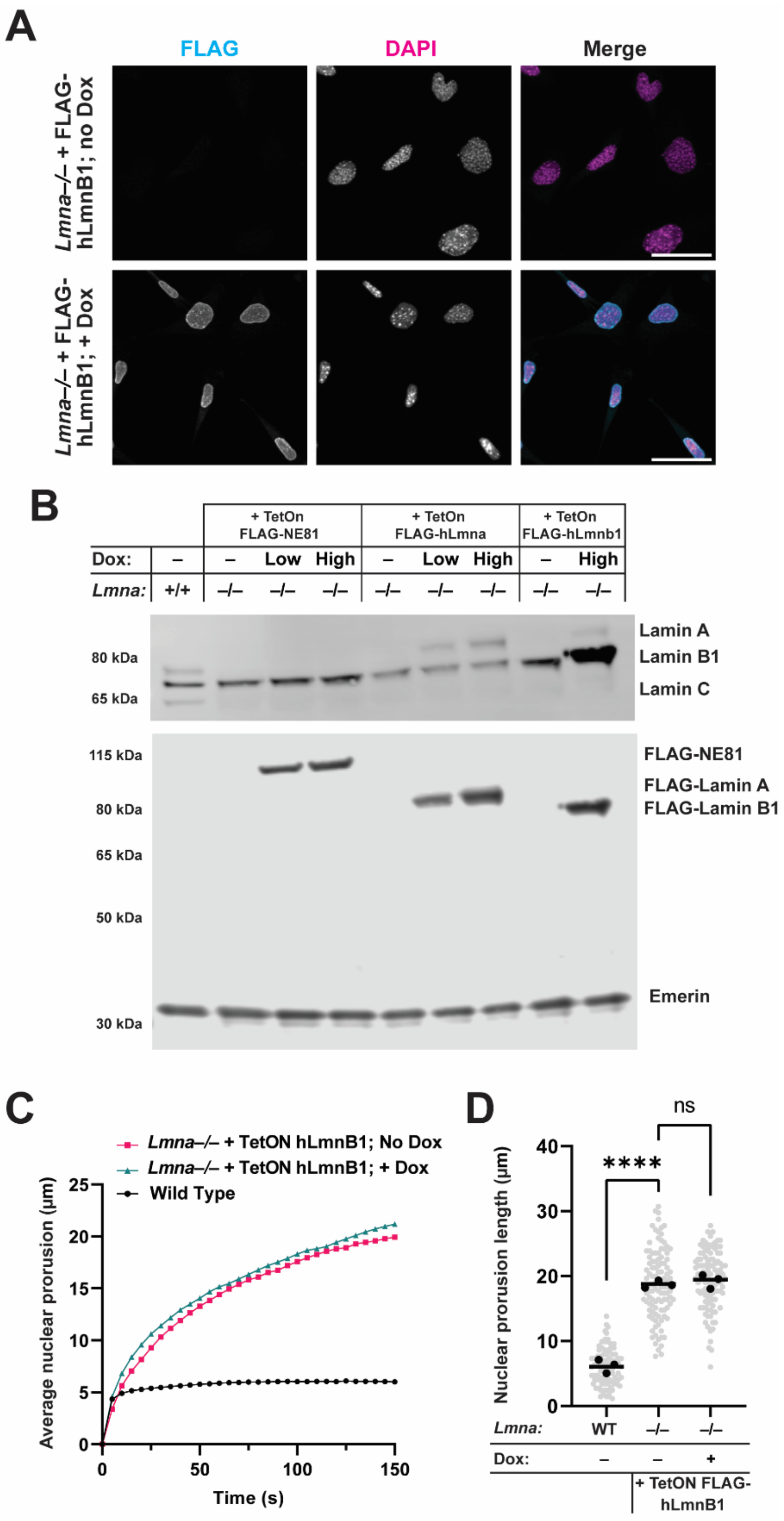


**Supplemental Figure 3: Inducible expression of FLAG-Lamin B1 in *Lmna*$^{-/-}$ MEFs. (A)** Representative immunofluorescence images of *Lmna*$^{-/-}$ MEFs expressing FLAG-human Lamin B1. Expression was induced by addition of 100 ng/ml doxycycline to the media for 24 h prior to fixation and staining. (**B**) Immunoblot analysis of cells expressing FLAG-NE81, FLAG-hLmna, or FLAG-hLmnb1. Membranes were probed with anti-FLAG antibody to demonstrate the relative expression levels of the different exogenous constructs. Antibodies against Lamin A/C and Lamin B1 were used to detect both endogenous and exogenously expressed proteins. Note that antibody epitope recognition may differ between the endogenous mouse lamins and the ectopically expressed human Lamin A and Lamin B1. Emerin staining was used as a loading control. (**C**) Quantification of average nuclear protrusion over time, using the same approach as in Figure 3. (**D**) Quantification of nuclear protrusion lengths 120 s after cells entered micropipette pocket, using the same settings as in Figure 3. **** $p < 0.001$; ns: not statistically significant. One-way ANOVA with Tukey's multiple comparison test. 90-110 cells were scored for each condition/treatment.

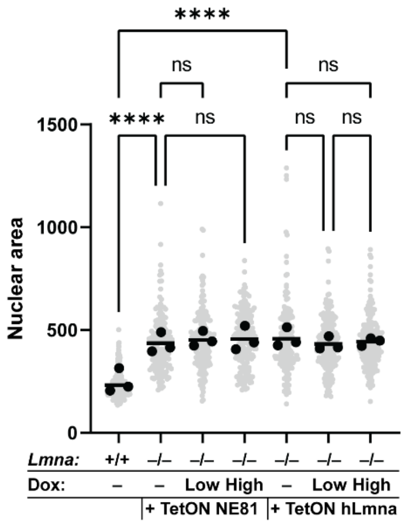


**Supplemental Figure 4: Nuclear cross-sectional area does not change upon expression of NE81 or Lamin A.** Nuclear cross-sectional (i.e., projected) areas of the same cells used in the quantification for Figure 4 were calculated from images segmented based on the DAPI staining for DNA. Area measurements are given in µm$^2$. **** $p$, < 0.001; ns: not statistically significant. Cell numbers are the same as in Fig. 4. One-way ANOVA with Tukey's multiple comparison test.

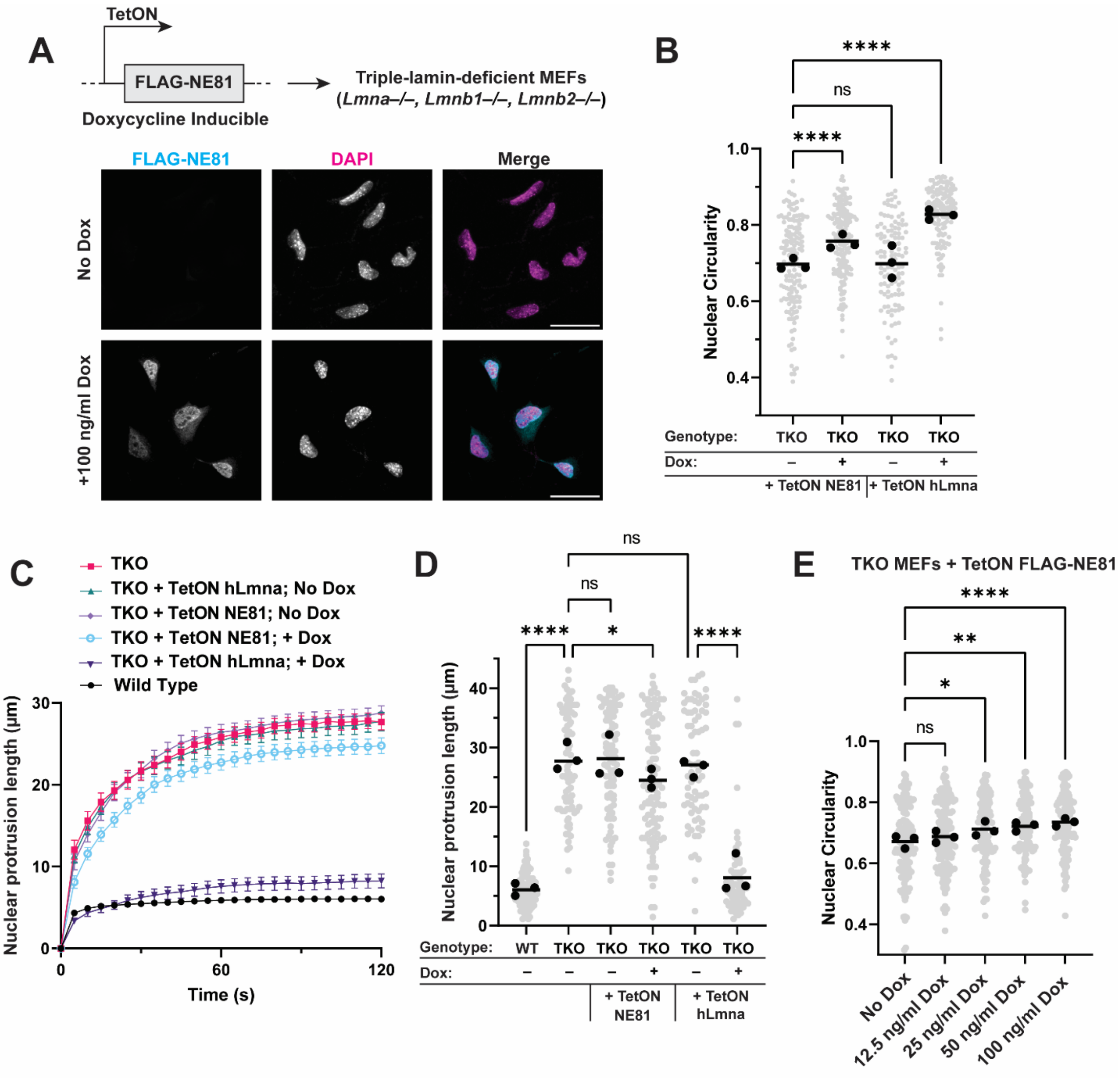


**Supplemental Figure 5: NE81 expression partially rescues morphological and mechanical defects in TKO MEFs that lack all lamins**. (**A**) Doxycycline-inducible NE81 expression in Triple-Lamin Knockout (*Lmna*$^{-/-}$, *Lmnb1*$^{-/-}$, *Lmnb2*$^{-/-}$) or TKO MEFs. Representative immunofluorescence images are shown. (**B**) Nuclear circularity analysis performed on cells in the presence or absence of doxycycline as in Figure 2. (**C**) Nuclear protrusion lengths over time following micropipette aspiration as in Figure 3. (**D**) Quantification of nuclear protrusion lengths 100 s after start of aspiration as in Figure 3. (**E**) Nuclear circularity of TKO MEFs expressing dox-inducible FLAG-NE81 with increasing dox concentrations added to the media does not show a dose-dependent response. "Plus dox" conditions in other figure panels refer to cells treated with 100 ng/ml doxycycline. * $p < 0.05$; ** $p < 0.01$; **** $p < 0.001$; ns, not statistically significant. One-way ANOVA with Tukey's multiple comparison test. For circularity measurements, 200-250 cells were scored per condition. For micropipette aspiration, 50-80 nuclei were measured per condition.
